# Curvature-based schedule ordering is preserved in homogeneous and heterogeneous tumors

**DOI:** 10.64898/2026.11.08.765156

**Authors:** A Hyeon Song, Jeffrey West

**Affiliations:** Department of Mathematics, University of Florida; Integrated Mathematical Oncology Department, H. Lee Moffitt Cancer Center & Research Institute

## Abstract

During treatment, dosing protocols with identical total dose may favor one schedule over another depending on how exposure is distributed over time. In homogeneous cell populations, Jensen’s inequality has been shown to predict whether continuous or intermittent high-dose followed by low-dose is favored, based on the curvature of the concentration-effect function. Scheduling ordering can be performed based on curvature only: concave functions favor continuous dosing while convex functions favor intermittent dosing schedules with a high time-variance. However, tumors exhibit heterogeneous drug sensitivities, and treatment preferentially eliminates drug-sensitive cells, progressively shifting the surviving population toward greater drug resistance. Whether this treatment-induced selection can reverse curvature-based schedule ordering remains unclear. We develop a mathematical framework in which phenotypes share a common concentration-effect function but differ by fixed, nonnegative sensitivity scales. We prove that the ordering of equal-mean schedules is administrated solely by the scalar cumulative treatment effect, that is, by the curvature of the effect function alone. Consequently, baseline heterogeneity and treatment-induced selection can alter the magnitude of treatment response and the composition of surviving cells, but cannot reverse the ordering of equal-mean schedules predicted by the curvature of the single-cell concentration-effect function. This preservation holds across different temporal dosing profiles and repeated treatment cycles. Our work uncovers a general principle that curvature-based schedule ordering is preserved in heterogeneous tumors despite treatment-induced selection toward resistance.

## Introduction

In cancer pharmacotherapy, mathematical modeling is increasingly being used to investigate and inform treatment dosing strategies.^12^ In practice, dosing strategies are often determined by combining a drug phar-macodynamic model of response to a range of dose concentrations together with a model of pharmacokinetic decay of drug through clearance^3^. Extensions of this framework consider intratumoral heterogeneity in drug sensitivity^4–6^. The objective function may vary, but typically aims to minimize the final tumor size, subject to some tolerability considerations of the maximum instantaneous or cumulative drug exposure. For most drugs, the current clinical practice uses a maximum tolerated dose (MTD) paradigm for dose-finding, which emerged from early pioneering work on chemotherapies for the treatment of leukemias^7, 8^. MTD dosing is used in phase I clinical trials, and subsequently typically only adjusted in the event of toxicity, interference with combination treatments, or tumor progression^9^. However, if tumor eradication is not achieved, relapse occurs due to the evolution of drug resistance^10^. Furthermore, MTD can lead to unnecessary toxicity and adverse events, which alternative therapeutic strategies could avoid^11^.

Pulsed administration of dosing has been proposed as a method for suppressing resistant cells with lower drug-sensitivity, by increasing the dose at lower frequencies^12, 13^. The efficacy of this approach may be partially supported by a curvature-effect, whereby the shape of the dose response function determines whether continuous or pulsed administration is favored^14^. Curvature-based treatment relies on the simple graphical principle known as Jensen’s inequality of the input (dose concentration) and output (tumor kill) functional relationship. If the dose response is convex, then pulsed administration is favored; if concave, then continuous administration.

Jensen’s inequality defines the relationship between function evaluated at the average input, compared to the average of the function evaluated at many unique inputs. For example, a dosing strategy with a highly varied series of doses (e.g. 200mg, 250mg, 180mg, …) can be compared to the dosing schedule of equivalent average dose (210mg). If the tumor kill function is convex, the varied dosing schedule is alwaysmathematicallyl superior to the continuous therapy equivalent, subject to the assumption that each dose provides an independent and additive effect. By Jensen’s inequality and the concepts of convexity and concavity, we can mathematically investigate whether differences in dosing schedule alone can generate therapeutic benefit, even when the mean exposure remains unchanged. This curvature-based rule provides a simple principle for comparing even (low variance, continuous schedules) and uneven dosing (high variance, intermittent) at equal mean exposure. The rule has been recently applied to link time-varying exposure with nonlinear treatment response^15, 16^. This curvature framework is particularly relevant to anticancer therapies, whose dose-response curves are typically sigmoidal or otherwise nonlinear rather than linear. Though dose response can be visualized on a variety of metrics such as cell survival^17^ or GR-curves^18^, the important metric of curvature is that of the functional relationship between dose and tumor kill rate (preserving independence and additivity). Previous work has shown that gains in tumor kill is possible without altering the total dose delivered, by leveraging knowledge of dose response curvature^19^.

Intermittent, pulsed administration may include alternative benefits. For example, periodic holidays from drug-dosing has been shown to resensitize the tumor population to subsequent drug doses in targeted therapies^19, 20^. This effect may be magnified in tumors with a strong cost of resistance^21, 22^ or observed drug-dependence^20, 23^.

### Intercellular heterogeneity alters drug-sensitivity

Tumors are heterogeneous^24, 25^. Cells vary continuously in drug sensitivity, and this baseline heterogeneity can cause the population-level dose-response relationship to differ substantially from the response of any single cell^26^. Treatment also does not act on a static population because more sensitive phenotypes are preferentially eliminated, the composition of the surviving population shifts over time (cycles) towards resistance^5, 27^. This compositional shift is often assumed to arise from pre-existing phenotypes rather than any treatment-induced adaptation or phenotypic transition^6, 10^. It is therefore unclear whether the curvature-based schedule ordering derived for a homogeneous population is preserved once tumor response is averaged over a heterogeneous distribution.

Multiple studies have shown, both in vitro^28–31^ and in vivo^32, 33^ that chemotherapy dose-response curves depend not only on a drug’s cell-cycle specificity but also on the duration of drug exposure. Such response are conventionally described by fitting a sigmoidal Hill function^30^. While useful for comparative potency comparison and for predicting clinical response, this approach is a statistical fit: it does not explain or predict how the properties of a drug or of the tumor cells shape the form of the dose-response curve, and it does not naturally accommodate temporal features of exposure^27^. In radiation oncology, for example, mathematical frameworks typically employ a log-survival model that decreases as a linear-quadratic function of dose, yet, this is a response form is specialized to a particular treatment modality^34, 35^.

Additionally, more recent models move beyond discrete sensitivity/resistance^36^ by describing drug sensitivity as a continuous distribution. The masking that occurs when averaging sensitivity across the cell population can distort apparent curvature: a convex, ultrasensitive cell-level response can, once integrated over a sensitivity distribution, yield an aggregate response that appears sublinear. Thus this apparent concavity would predict that even treatment is optimal yet concentrated, uneven dosing is often observed to remain favored clinically^6^. However, to solve this question, curvature-based analyzes and heterogeneous population mathematical models have recently been addressed, and they were developed for particular dose-response forms, specific survival functions, or specific cancers. It still remains unclear whether the homogeneous-population schedule ordering is preserved for an arbitrary baseline sensitivity distribution when under drug-induced selection pressure.

### Curvature-based scheduling ordering in homogeneous or heterogeneous tumors

In this paper, we investigate whether curvature-based schedule ordering derived for homogeneous populations remains preserved under heterogeneous drug sensitivity and treatment-induced selection. To address this question, we consider a mathematical model of heterogeneous tumors that captures selection among pre-existing phenotypes with different drug sensitivities. We begin by establishing the curvature-based scheduling rule using Jensen’s inequality (Fig.1). We then extend this framework from homogeneous populations to discrete and continuous heterogeneous sensitivity distributions, introducing the cumulative treatment effect as a common quantity linking temporal drug exposure to population survival (Fig.2). Next, we examine whether this ordering is preserved across different temporal dosing profiles, including step, sinusoidal, and pharmacokinetic schedules (Fig.3). We further investigate how treatment-induced selection changes the sensitivity distribution of surviving cells over repeated treatment cycles (Fig. 4). Finally, we examine the robustness of curvature-based schedule ordering across different baseline sensitivity distributions, including lognormal, gamma, and Weibull distributions (Fig.5). Together, these analyses establish a general framework for understanding how curvature-based schedule ordering is preserved across heterogeneous tumors despite treatment-induced selection toward resistance.

**Figure 1.**
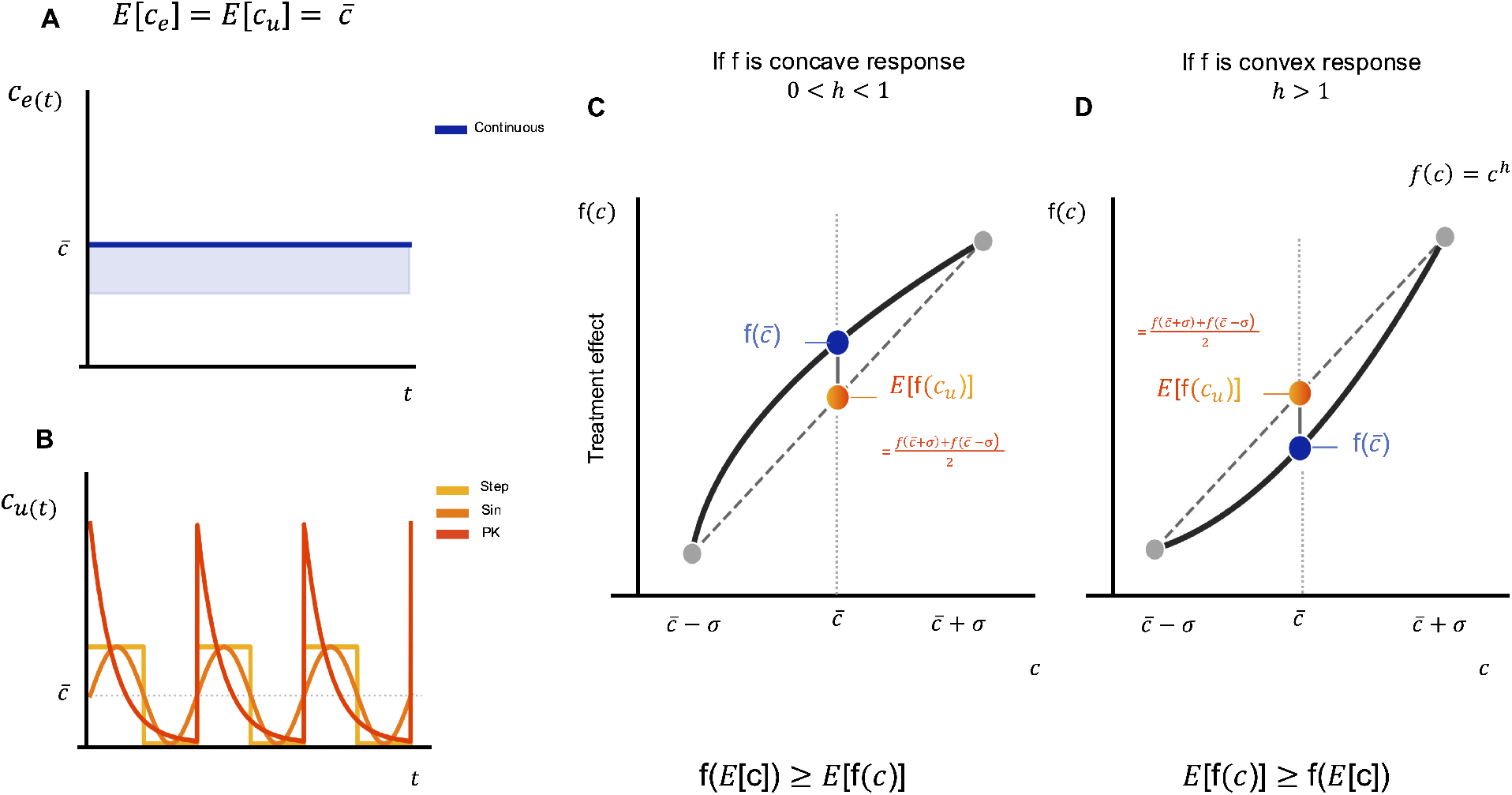
Jensen inequality determines the effect of temporal concentration variation under equal-mean treatment schedules. A, Even schedule 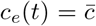. B, Uneven step, sinusoidal, and Pharmacokinetic (PK) schedules with the same temporal mean, 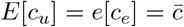. C, for a concave response *f* (*c*)= *c*^*h*^, 0 <*h*< 1, Jensen’s inequality gives 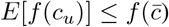, favoring the even schedule in cumulative treatment effect. D, for a convex function, *h*> 1, 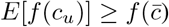, favoring the uneven schedule. For *h* = 1, the schedules are equivalent because *f* (*c*) is linear.

**Figure 2.**
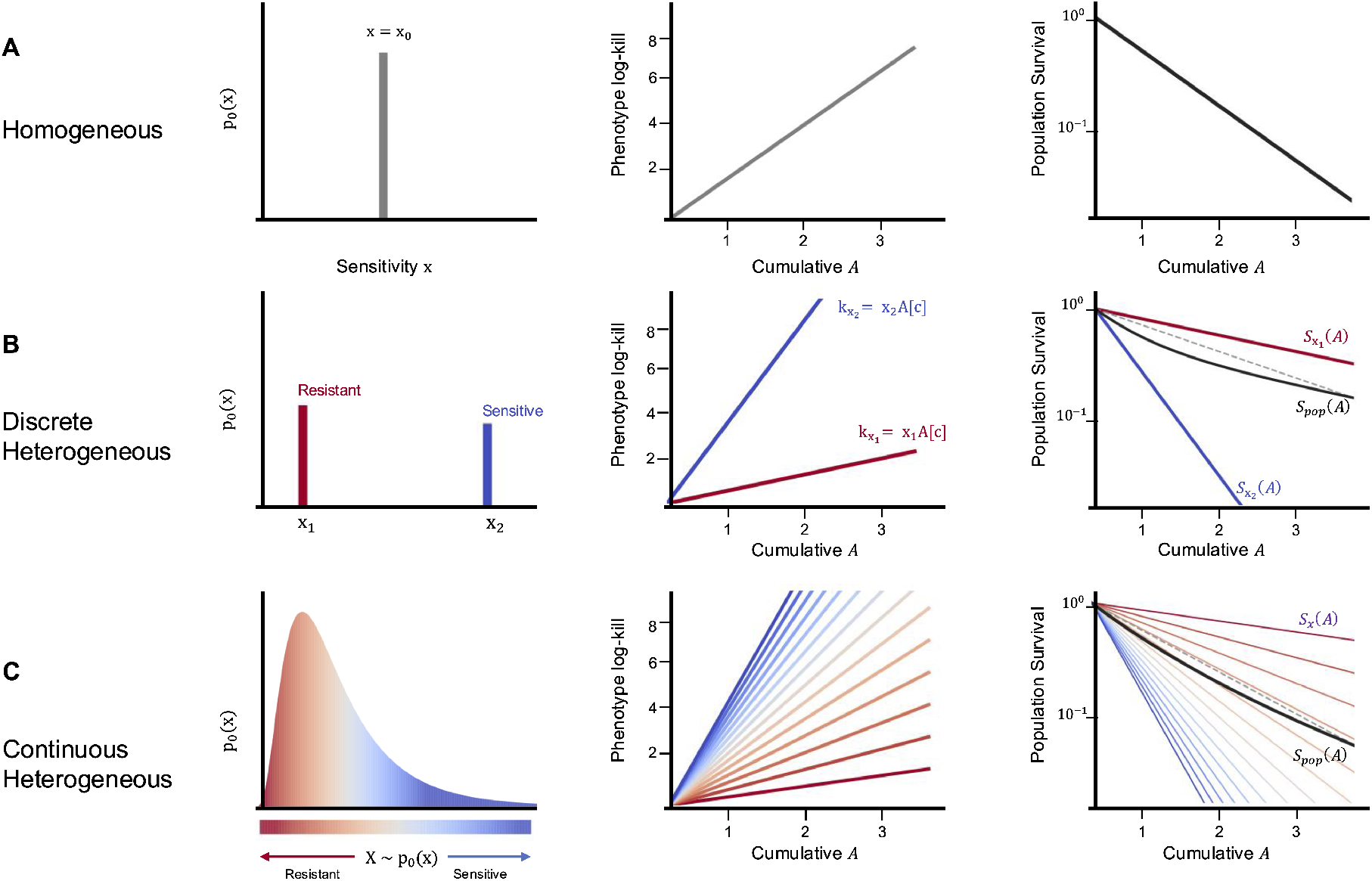
Rows: homogeneous, discrete, and continuous baseline sensitivity. Column 1: Baseline distribution *p*_0_(*x*). Column 2: Phenotype log-kill *K* = *xA*, each clone follows the same linear law in A, with slope set by its sensitivity *x*. Column 3: Population remaining fraction *S*(*A*). In the homogeneous cases, 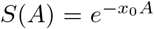. With discrete or continuous heterogeneity, 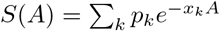 or *S*(*A*)= *E*_*p*0_ [*e*^−*XA*^], so the aggregate curve can depart from a single-rate exponential (dashed chord), yet every phenotype still obeys *S*_*x*_ = *e*^−*xA*^ and every schedule affects the population only through the scalar *A*[*c*].

**Figure 3.**
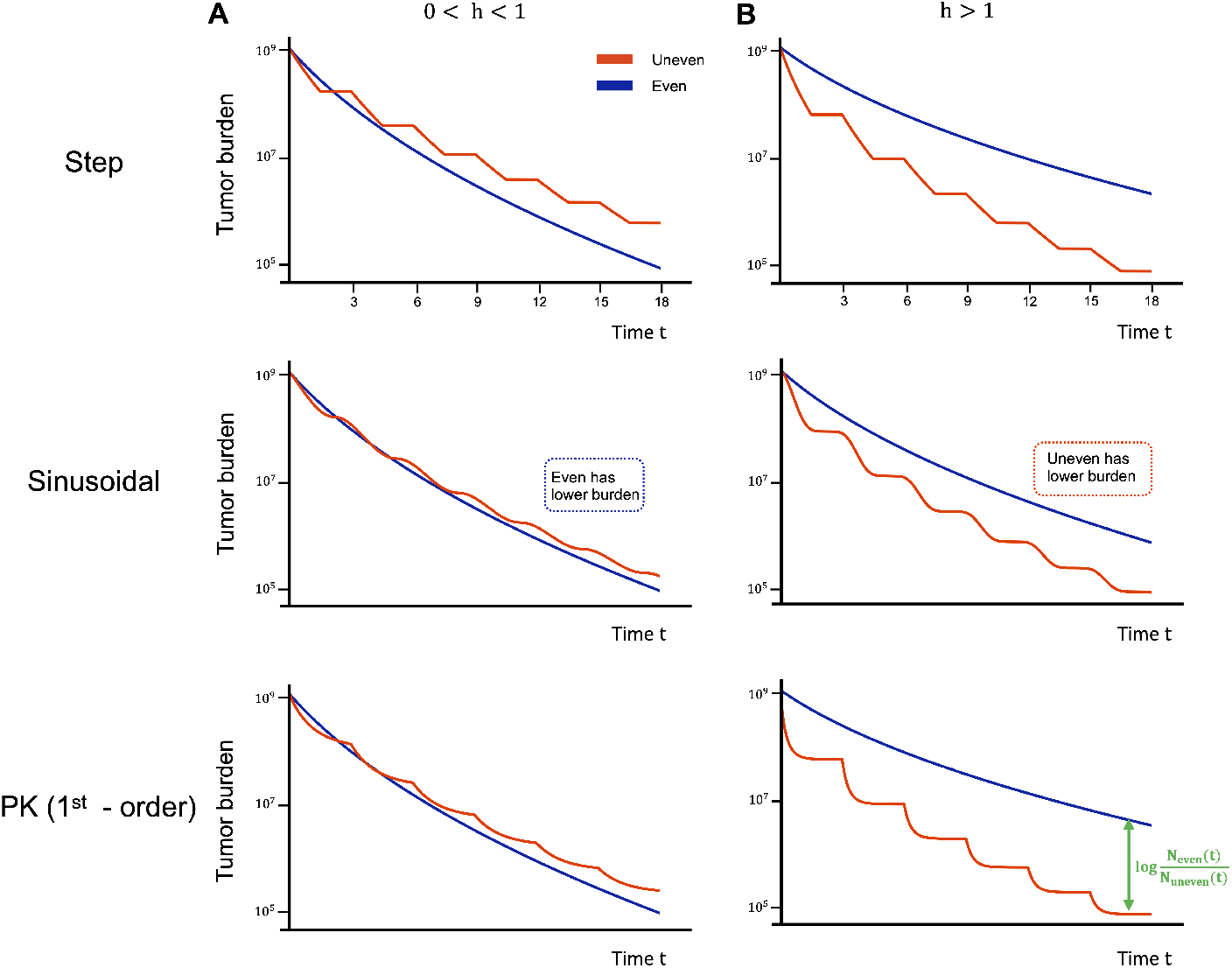
Tumor burden under even (blue) and uneven (orange) equal-mean schedules is shown for step, sinusoidal, and first-order-pharmacokinetic decay concentration profiles. A. For a concave concentration-effect function, (*h* = 0.5), the even schedule has the larger cumulative treatment effect, *A*[*c*_*u*_] ≤ *A*[*c*_*e*_], and produces the lower final tumor burden. B, For a convex response, (*h* = 2), the ordering reverses, *A*[*c*_*u*_] ≥ *A*[*c*_*e*_], and the uneven schedule produces the lower final burden. Although the temporal trajectories depend on the concentration profile, the final schedule ordering is determined by the ordering of *A*[*c*]. Simulations use (*T* = 18), 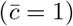, and a lognormal baseline sensitivity distribution.

**Figure 4.**
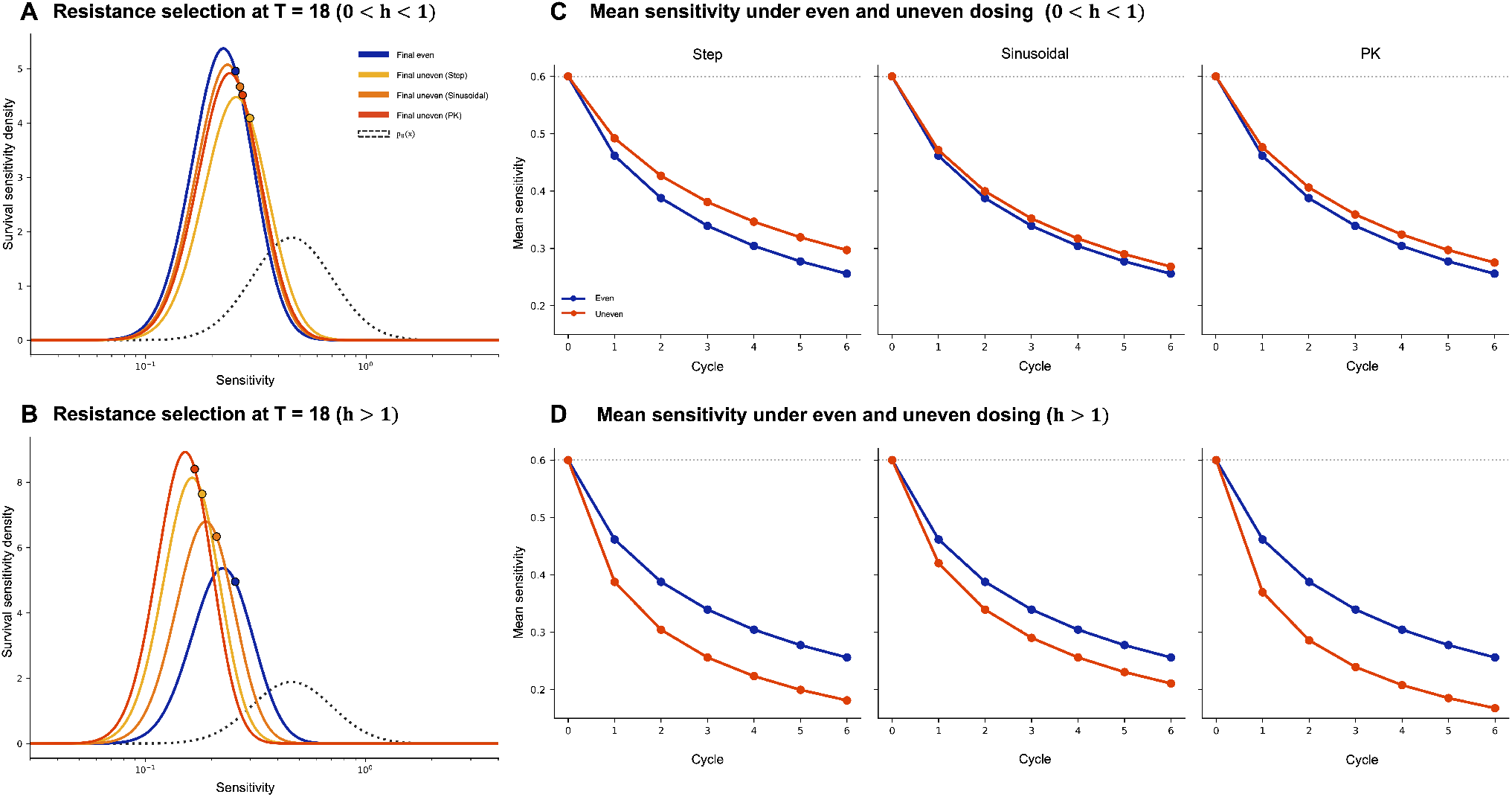
A-B, Final survivor sensitivity distributions after even and uneven dosing for concave and convex response. C-D, Mean sensitivity of surviving cells across treatment cycles for step, sinusoidal, and PK schedules. Larger cumulative treatment effect *A*[*c*] produces stronger reduction of sensitive phenotypes and a greater shift toward resistant survivors.

**Figure 5.**
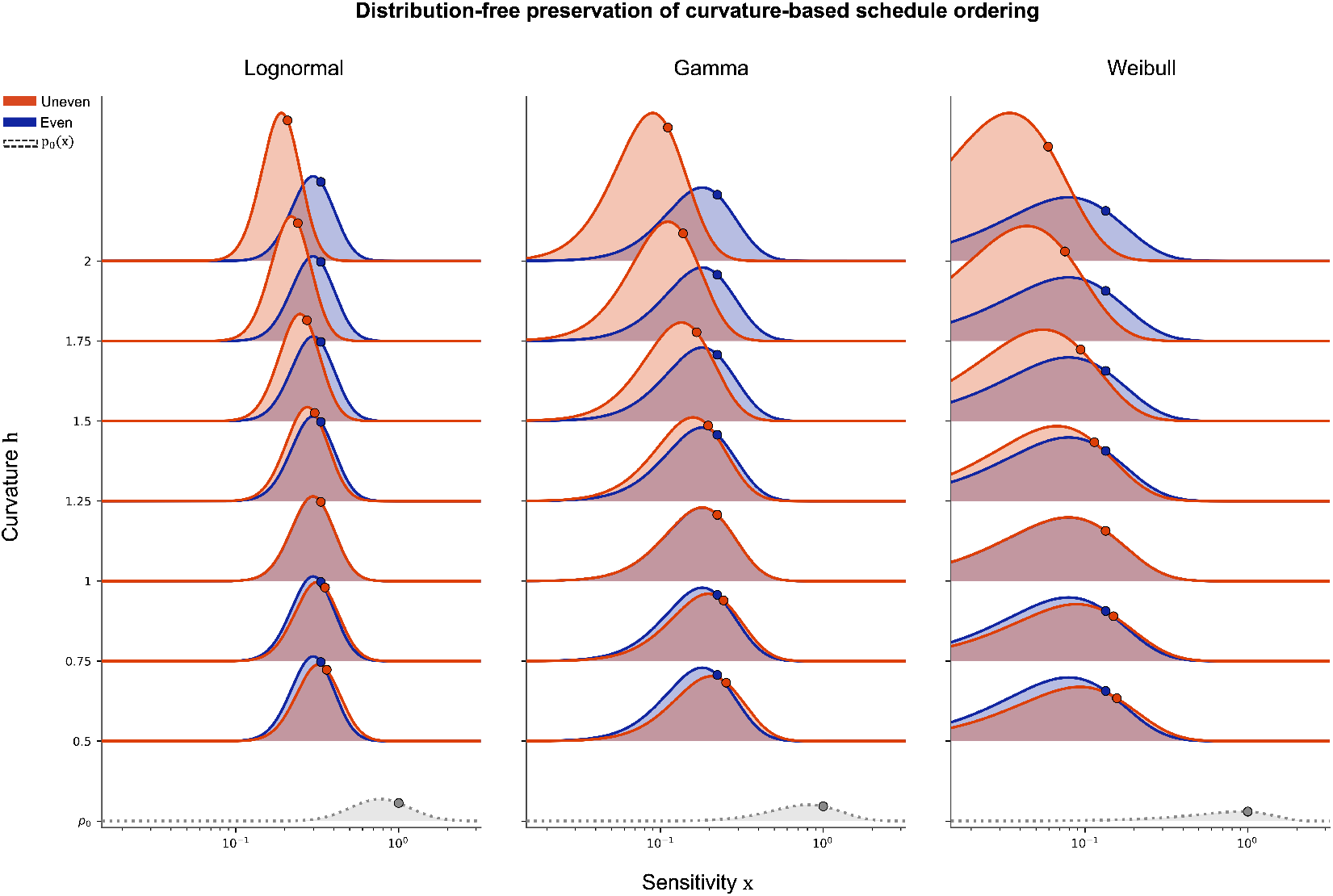
Final survivor sensitivity distributions are shown after even (blue) and uneven (orange) equal-mean dosing for lognormal, gamma, and Weibull sensitivity distributions *p*_0_(*x*) (gray dotted curves). Rows vary the curvature parameter *h* of the concentration-effect function *f* (*c*) = *c*^*h*^. All baseline distributions are matched to have men (*E*[*x*] = 1) and variance (Var(*x*) ≈ 0.1929). For convex response (*h* > 1), uneven dosing generates larger cumulative treatment effect and shifts the surviving population further toward lower sensitivity. For concave response (0 <*h* < 1), the ordering reverses, and even dosing produces the stronger shift toward resistance survivors. At *h* = 1, the two schedules are equivalent.

## Methods

### Jensen’s inequality to determine schedule ordering

We prove that for any baseline sensitivity distribution *p*_0_(*x*), the ordering of two equal-mean schedules is governed entirely by the scalar cumulative treatment effect *A*[*c*] and hence by the curvature of the cell-level response alone. Baseline heterogeneity rescales the final tumor burden and can alter the apparent curvature of the population response, but it cannot reverse which schedule minimizes that burden. Consider Jensen’s inequality, as a point of departure for setting up our curvature-based treatment scheduling framework. Jensen’s inequality compares the expected value (the average) of outcomes under two scenarios: zero input variance and high input variance. More precisely, for a function *q* and a varying quantity *u*, the average output differs from the output at the average input, with the direction fixed by the curvature of *q*:

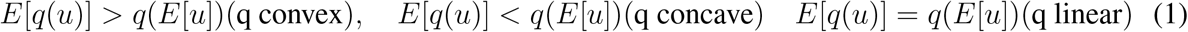

Equivalently, the curvature of *q* alone fixes the direction of the inequality: increasing the variance of the input *u* raises the average output above *q*(*E*[*u*]) when *q* is convex and lowers it when *q* is concave.^37, 38^ Therefore, the gap is

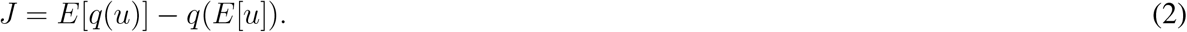

Without loss of generality, we consider the response to drug as our output function and the drug concentration as our input function. Consider a drug administered with an arbitrary dosing protocol resulting in a concentration, *u* := *c*(*t*), over time. The response of the tumor is given by *q*(*u*) := *f* (*c*), called the dose-response function. Our goal is to determine the schedule “ordering” (identify the dosing schedule which results in maximum tumor kill) for two dosing schedules.

First, we consider an arbitrary, temporally varying schedule, *c*_*u*_(*t*), with non-zero variance about the mean dose. By terminology, we refer to this as an “uneven” schedule because of its non-zero variance. For comparison, we consider whether this schedule outperforms or underperforms a second dosing schedule with identical mean, but zero variance. By terminology, we refer to this as an “even” treatment schedule 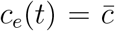 satisfying the equal mean constraint 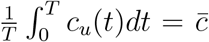. The even schedule has zero temporal variance, whereas the uneven schedule has positive temporal variance.

### Growth dynamics with homogeneous drug sensitivity

Let *n*(*x, t*) denote the abundance of tumor cells with fixed baseline drug sensitivity *x* ≥ 0 at time *t*. We assume that all phenotypes share the same concentration-effect function *f* (*c*), but differ through the non-negative drug-sensitivity scalar *x*. Under treatment concentration schedule *c*(*t*), phenotype *x* follows:

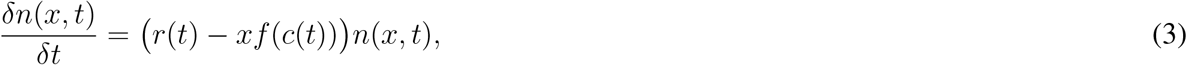

where *r*(*t*) is the treatment-independent net proliferation rate, *f* (*c*(*t*)) ≥ 0 is the common concentration-dependent treatment effect, and *x* is the fixed phenotype-specific sensitivity. Thus *xf* (*c*(*t*)) represents the treatment-induced loss rate for cells with sensitivity phenotype of *x*. Cell sensitivity, *x*, is fixed during treatment: the model describes selection among pre-existing phenotypes rather than treatment-induced phenotype switching or adaptation. Integrating Eq. 3 between a fixed treatment time interval, [0,*T*] gives the following separable solution:

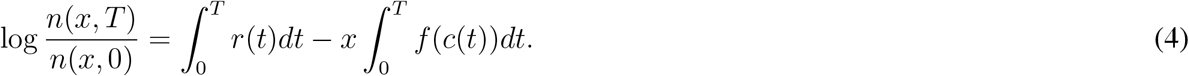

By notation, we define the cumulative treatment effect 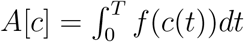 and the treatment-independent growth factor 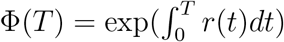. Then Eq. 4 can be written:

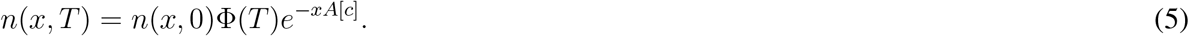

Dose response functions are commonly written as a survival fraction relative to untreated growth. Untreated growth is given by *n*_untreated_(*x, T*) = *n*(*x*, 0)Φ(*T*), and thus the fraction of surviving cells subject to a dose of *c* reduces to:

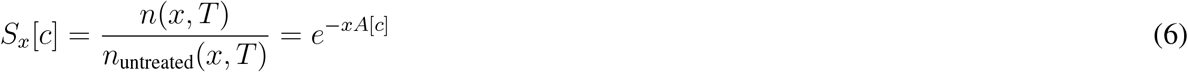

## Intratumoral heterogeneity in sensitivity to drug

However, not all cells have equivalent sensitivity to a given dose. Let *X* denote a nonnegative random variable representing drug sensitivity, with baseline probability density *p*_0_(*X* = *x*). The initial tumor population is given by

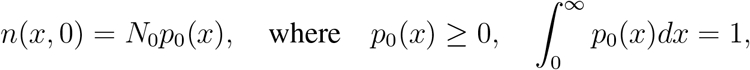

Substitution into (Eq.5) gives

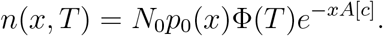

The total tumor burden is therefore

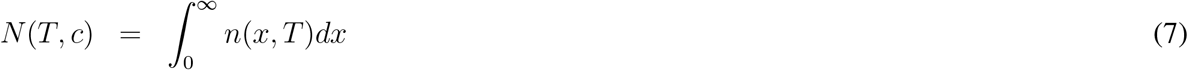

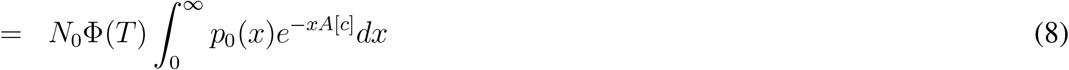

Thus, all temporal information about the treatment schedule’s efficacy is compressed into the single scalar *A*[*c*], whereas baseline heterogeneity enters through the expectation over *X*.

The ordering preservation. Let *c*_1_, *c*_2_ be two schedules. Suppose *X* ≥ 0. If

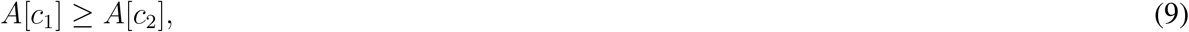

then

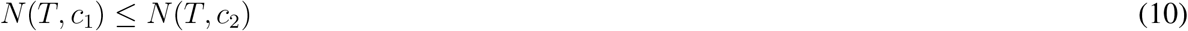

for every baseline-sensitivity distribution *p*_0_.

*Proof*. Since *x* ≥ 0, *A*[*c*_1_] ≥ *A*[*c*_2_] implies *xA*[*c*_1_] ≥ *xA*[*c*_2_] for ∀*x* ≥ 0. Because *e*^−*xA*^ decreases monotonically with *A*, ∀*x* ≥ 0. Multiplying by *p*_0_(*x*) ≥ 0, integrating over *x*, and multiplying by *N*_0_ Φ(*T*) > 0 gives *N* (*T, c*_1_) ≤ *N* (*T, c*_2_).

For any baseline distribution of non-negative drug sensitivities, a larger cumulative treatment effect produces a final tumor burden that is no greater than that produced by a smaller cumulative treatment effect. This ordering is independent of the shape of the baseline sensitivity distribution. See Section S1 for more information.

## Preservation of Jensen schedule ordering

As stated above, let the even schedule be 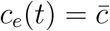 and let *c*_*u*_ be any non-constant schedule of equal mean, 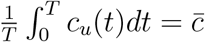. If *f* is concave, then by (Eqs.9-10):

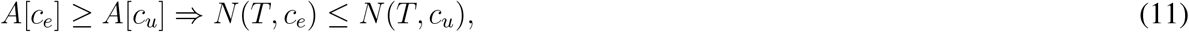

so, the even schedule yields the lower burden. If *f* is convex, the inequalities reverse and the uneven schedule yields the lower burden. Therefore,

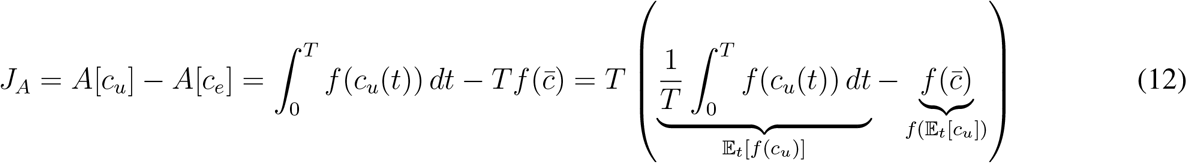

By Jensen’s inequality, concave concentration-effect functions favor even exposure, whereas convex functions favor uneven exposure under equal-mean schedules. Because of our focus on the curvature of the dose effect function, we consider a power-law concentration-effect function, *f* (*c*) = *c*^*h*^, that can allow for linear (*h* = 1), convex (*h*> 1), and concave (*h*< 1) shapes. Jensen’s inequality gives sgn(*J*_*A*_) = sgn(*h* −1). Thus, *J*_*A*_ < 0 for 0 < *h* < 1, *J*_*A*_ > 0 for *h* > 1. The corresponding difference in final tumor burden is defined as

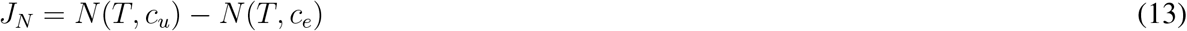

By the ordering preservation established in (Eqs.9-10), a larger cumulative treatment effect produces a final tumor burden that is no greater than that produced by a smaller cumulative treatment effect, regardless of the baseline sensitivity distribution. Consequently, when the baseline population contains cells with positive drug sensitivity, Therefore, sgn[*N* (*T, c*_*u*_) − *N* (*T, c*_*e*_)] = -sgn(*h* − 1). Hence the even schedule is favored for 0 <*h* < 1, whereas the uneven schedule is favored for *h*> 1, independently of the baseline sensitivity distribution *p*_0_.

Let 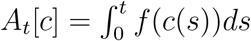 be for the cumulative effect up to time *t*. The baseline-sensitivity distribution among survivors is

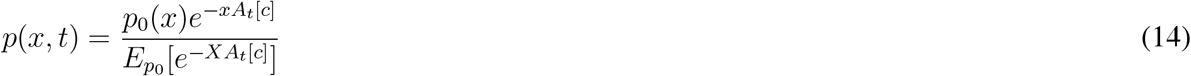

The mean sensitivity among survivors is

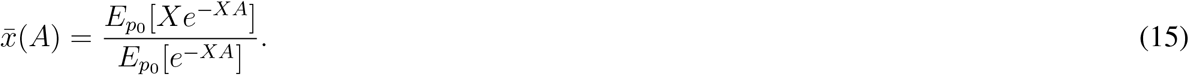

Differentiating with respect to cumulative treatment effect gives

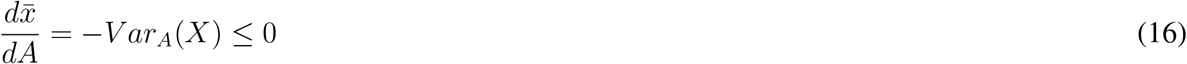

Therefore, increasing cumulative treatment exposure preferentially removes phenotypes with larger *x*. See Section S1 for more information. This approach can be used to describe discrete dosing, via summation:

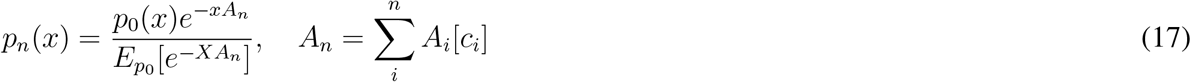

Partition the horizon into *n* successive cycles on intervals [*t* _*i*−1_, *t*_*i*_], and let 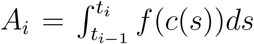 be the cumulative effect delivered in cycle *i*. Because the cumulative effect is additive over time, 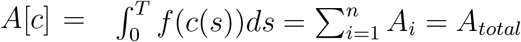, and the final burden depends on the schedule only through *A*_*total*_,

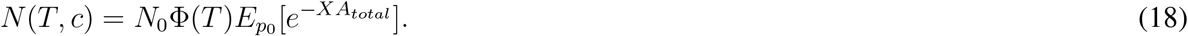

Thus, intermediate selection across subsequent treatment cycles does not create an additional dependence on a schedule’s time history. Final burden depends on the treatment trajectory only through the total cumulative effect *A*_*total*_. Consequently, selection across multiple cycles cannot reverse the ordering induced by *A*[*c*].

Since selection shifts survivors toward resistance, any *A*_*t*_[*c*] ≥ 0, the survivor distribution *p* is smaller than *p*_0_ in the monotone-likelihood-ratio order, and hence

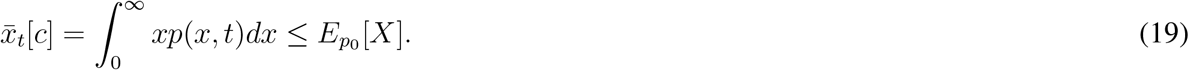

The survivor mean is strictly decreasing in cumulative exposure, so larger cumulative exposure produces a more resistant surviving population. Therefore, the schedule affects the burden only through the scalar *A*[*c*], and its ordering preserved for every *p*_0_.

### Relevant examples of dosing schedules

For practical motivation of the above proofs, we provide several examples of dosing schedules with intrinsic input temporal variance in dosing: step functions, sinusoidal functions, and pharmacokinetic decay functions (see Table 1). To prove the ordering across qualitatively distinct temporal profiles, we fix three schedules of equal mean 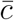.

**Table 1.** Representative equal-mean treatment schedules and their cumulative effects under the power-law concentration-effect function *f* (*c*)= *c*^*h*^.

| Schedule | Concentration profile $c(t)$ | Equal-mean condition | Cumulative effect $A_h[c]$ |
|---|---|---|---|
| Step | $c_{\mathrm{step}}(t) = \begin{cases} c_H, & 0 \le t < \theta T, \\ c_L, & \theta T \le t \le T \end{cases}$ | $\theta c_H + (1-\theta) c_L = \bar{c}$ | $T\left[\theta c_H^h + (1-\theta) c_L^h\right]$ |
| Sinusoidal | $c_{\sin}(t) = \bar{c} + a \sin\left(\frac{2\pi t}{T} + \phi\right)$ | $\frac{1}{T}\int_0^T c_{\sin}(t)\,dt = \bar{c}$ | $\int_0^T \left[\bar{c} + a \sin\left(\frac{2\pi t}{T} + \phi\right)\right]^h dt$ |
| PK | $c_{\mathrm{PK}}(t) = c_0 e^{-kt}$ | $\bar{c} = \frac{c_0}{kT}(1 - e^{-kT})$ | $\frac{c_0^h}{hk}(1 - e^{-hkT})$ |

## Results

### Curvature determines the ordering of equal-mean treatment schedules

We develop the framework by first isolating the effect of temporal variation in drug concentration before introducing tumor heterogeneity. We begin with an even treatment schedule, 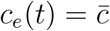, and compare it with temporally uneven schedules *c*_*u*_(*t*) constrained to have the same mean concentration over the treatment horizon. (Fig. 1A, B). The uneven schedules included step, sinusoidal, and first-order pharmacokinetic-like profiles, all satisfying 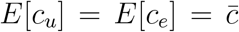. Despite having same mean concentration, these schedules need not produce the same treatment effect when the concentration-effect function *f* is nonlinear. In the model, the temporal contribution of a schedule is summarized by the cumulative treatment effect 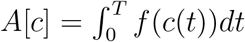. Thus, schedules with the same mean exposure can differ in *A*[*c*] solely because of how concentration is distributed over time.

Jensen’s inequality determines the direction of this difference. For a concave concentration-effect function, 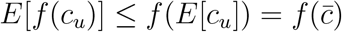, so that *A*[*c*_*u*_] ≤ *A*[*c*_*e*_], so temporal variation decreases cumulative treatment, favoring the even schedule. (Fig. 1C). For a convex concentration-effect function, the inequality reverses, giving *A*[*c*_*u*_] ≥ *A*[*c*_*e*_], so temporal variation increases the cumulative treatment effect (Fig. 1D). When *f* is linear the two schedules are equivalent. Thus, for equal-mean schedules, the direction of schedule ordering is preserved by the curvature of the cell-level concentration-effect function rather than by the specific temporal profile. In a homogeneous population, the curvature of the concentration-effect function alone determines whether temporal variation increases or decreases cumulative treatment effect.

### Heterogeneous phenotypes share a common dependence on the cumulative treatment effect

The cumulative treatment effect *A*[*c*] provides a common treatment coordinate across phenotypes with different baseline sensitivities. From Eqs.7, the phenotype-specific response relative to untreated growth is 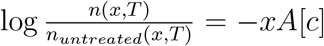. Defining the remaining fraction, Eqs.7 therefore give −*logS*_*x*_[*c*] = *xA*[*c*], *S*_*x*_[*c*] = *e*^−*xA*[*c*]^. Thus, every phenotype responds to the same schedule-dependent quantity *A*[*c*], while the sensitivity *x* determines only the strength of that response. On a log-survival scale, phenotype response are therefore linear in *A*[*c*], with slope *x* (Fig.2).

We first consider a familiar population, homogeneous, *p*_0_(*x*) = *ω*(*x* − *x*_0_), in which all cells have sensitivity *x*_0_ (Fig. 2A). The population then follows the single phenotype response, 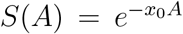, and log-kill increases linearly with *A* at rate *x*_0_. Next, consider a discrete heterogeneous population containing phenotypes with different sensitivities *x*_1_ and *x*_2_ (Fig.2B). Each phenotype retains the same linear dependence on *A*, but with a different slope: more sensitive phenotypes are reduced more rapidly, whereas less sensitive phenotypes as treatment accumulates.

More sensitive phenotypes therefore decline more rapidly with increasing *A*, whereas less sensitive phenotypes persist. In other words, as treatment accumulates, the contribution of the more sensitive phenotypes decays more rapidly, causing population survival to become progressively dominated by the resistant phenotype.

The same structure extends to a continuous sensitivity distribution *p*_0_(*x*) (Fig.2C). For *X* ∼ *p*_0_, averaging phenotype-specific survival over the population gives *S*(*A*[*c*]) = *E*_*p*0_ [*e*^−*XA*[*c*]^], and therefore *N* (*T, c*) = *N*_0_Φ(*T*)*E*_*p*0_ [*e*^−*XA*[*c*]^]. Heterogeneity therefore changes the mapping between cumulative treatment effect and population burden: a homogeneous population follows a single exponential, whereas a heterogeneous population is a mixture of phenotype-specific exponential. Importantly, however, heterogeneity introduces no additional dependence on the temporal treatment schedule.

Previous work^6^ illustrated a masking effect that occurs when observing only the tumor scale. In Fig.2, Column 3, the black curve shows this aggregate population response. Whereas the homogeneous population follows a single exponential response, heterogeneous populations progressively deviate from the corresponding single-rate exponential reference shown by the dashed gray line. Yet, this change in the population-level response does not introduce a new dependence on treatment schedule. Heterogeneity changes the mapping from *A* to population survival, as illustrated by the black curves, but the temporal schedule still enters every phenotype and the aggregate population only through the same scalar *A*[*c*].

### Cumulative treatment effect preserves schedule ordering across heterogeneous tumors

Next, we asked whether the ordering established through *A*[*c*] predicts tumor response when treatment is applied over time to a heterogeneous population. We compare even and uneven schedules with the same mean concentration using the step, sinusoidal, and first-order pharmacokinetic (PK) profiles introduced in (Table.1).

Although these schedules produce distinct tumor-burden trajectories, their final ordering remained determined by the cumulative treatment effect *A*[*c*] (Fig.3). For a concave concentration-effect function, (0 <*h* < 1), Jensen’s inequality gives *A*[*c*_*u*_] ≤ *A*[*c*_*e*_]. Because population burden is a decreasing function of *A*, the even schedule consequently produced the lower final tumor burden for each of the three temporal profiles (Fig. 3A). Conversely, for a convex response, (*h* > 1), *A*[*c*_*u*_] ≥ *A*[*c*_*e*_], and the uneven schedule produced the lower final burden (Fig. 3B). Thus, step, sinusoidal, and first-order PK decay schedules differed substantially in their temporal dynamics, but the direction of their final treatment advantage was preserved. Therefore, the specific temporal shape of the schedule determines the trajectory of response, whereas response curvature determines which equal-mean schedule ultimately produces the lower tumor burden.

### Treatment-induced selection shifts survivors toward resistance without reversing schedule ordering

Preservation of final tumor-burden ordering does not imply that treatment leaves the heterogeneous population unchanged. As treatment progressed, not only tumor burden but also the composition of the surviving population changed through Darwinian selection. More sensitive phenotypes were preferentially eliminated by treatment, causing the surviving population to become progressively enriched for less sensitive phenotypes over repeated treatment cycles. Consequently, the mean sensitivity of the surviving population decreased over time (Fig.4C, D).

The strength of this selection was determined not by the schedule type itself, but by the cumulative treatment effect generated by that schedule. For a concave response, the even schedule produced the larger cumulative treatment effect and therefore more strongly depleted sensitive phenotypes, resulting in a greater reduction in the mean sensitivity of survivors (Fig. 4A). In contrast, for a convex response, the uneven schedule produced the larger cumulative treatment effect and consequently induced a stronger shift toward resistant phenotypes (Fig.4B).

Under even dosing, the survivor distribution shifted smoothly toward lower sensitivity because cumulative treatment effect accumulated continuously at a constant rate. Under PK dosing, selection proceeded non-uniformly within each cycle, rapid shifts followed drug administration, whereas slower changes occurred as concentration decayed. These cycle-dependent changes were modest for convex response, where high-concentration exposure disproportionately increases treatment effect and concentrates selection immediately after each dose.

Thus, the schedule producing the larger cumulative treatment effect simultaneously reduced tumor burden more strongly and exerted greater selective pressure on sensitive phenotypes, thereby more strongly re-weighting the surviving population toward resistance. Importantly, this treatment-induced selection did not reverse the schedule ordering observed above. Although population composition continued to change throughout treatment, the schedule that produced the lower tumor burden remained determined by the curvature of the concentration–effect function. The schedule that produces the larger cumulative treatment effect also produces stronger selection toward resistant survivors. Selection changes who remains, but does not reverse which schedule produces the lower burden.

### Curvature-based schedule ordering is preserved across sensitivity distributions

Next, to examine whether the ordering depends on the assumed form of baseline heterogeneity, we compared lognormal, gamma, and Weibull sensitivity distributions matched in their mean and variance. Matching the first two moments allowed the effect of distributional shape to be varied without simultaneously changing the average sensitivity or overall degree of heterogeneity. Across all three distributions as shown in (Fig 5), changing the curvature parameter *h* altered the relative cumulative effect of even and uneven dosing in the direction predicted by Jensen’s inequality. For a concave response (0 < *h* < 1), even dosing produced a larger cumulative treatment effect and a stronger shift of the surviving population toward lower sensitivity. As *h* approached 1, the difference between the two schedules diminished, and *h* = 1 the survivor distributions overlapped, consistent with the equivalence of equal-mean schedules under a linear concentration-effect relationship. For a convex response (*h*> 1), this ordering reversed, with uneven dosing producing the stronger shift toward resistant survivors. Although the shape of the post-treatment sensitivity distribution across baseline distributions, the direction of schedule ordering was preserved across distributional forms at each value of *h*.

## Discussion

Therapeutic response in heterogeneous tumors reflects the intertwined effects of temporal drug exposure and treatment-induced selection. In vivo, drug concentrations vary continuously after administration as a consequence of distribution, metabolism, and elimination, while conventional in vitro drug-response assays commonly expose cells to fixed concentrations. Experimental systems that reproduce dynamic, in vivo-like pharmacokinetic profiles have shown that altering the temporal pattern of exposure can change treatment efficacy, demonstrating that drug response depends on more than a single nominal concentration or peak exposure.^39^ At the same time, treatment acts selectively on heterogeneous populations, preferentially depleting more sensitive cells and thereby reweighting the surviving population toward less sensitive phenotypes. High-complexity lineage tracing has directly shown that drug treatment can select rare pre-existing resistant clones,^40^ while longitudinal barcoding experiments have further demonstrated that treatment dose, duration, sequence, and drug holidays can jointly alter both population response and clonal composition.^41^ These processes make population-level treatment outcomes difficult to attribute mechanistically. Consequently, an observed difference between the treatment schedules combines effects arising from the temporal concentration profile with effects arising from different survival across heterogeneous phenotypes. Our model separates these contributions by first mapping the concentration trajectory *c*(*t*) to a schedule-dependent cumulative treatment effect *A*[*c*], and then translating this common treatment coordinate into population response through phenotype-specific sensitivities *x*.

Our mechanistic analysis of this decomposition shows that temporal scheduling and population heterogeneity play qualitatively different roles in treatment response. For equal-mean schedules, the direction of the treatment advantage is determined by the curvature of the concentration-effect function through the cumulative effect *A*[*c*] : concave responses favor even exposure, while convex responses favor uneven exposure. This ordering is preserved across heterogeneous populations because all phenotypes experience the same schedule-dependent coordinate *A*[*c*], differing only in the strength of their response through the sensitivity scale *x*. Heterogeneity therefore changes the magnitude and shape of the aggregate population response, but not the direction of schedule ordering. Importantly, this remains even as treatment progressively reweights the surviving population toward less sensitive phenotypes. More sensitive cells are preferentially depleted, so the survivor distribution becomes increasingly resistant over time, yet this dynamic selection does not introduce an additional schedule dependence capable of reversing the ordering set by *A*[*c*]. Thus, curvature determines which schedule is favored, whereas heterogeneity and selection determine how strongly the population responds and which phenotypes remain.

This preservation extends beyond the presence of heterogeneity itself to the form of the baseline sensitivity distribution. Across distinct baseline distributions, the magnitude and composition of the surviving population can differ substantially, while the relative ordering of equal-mean schedules remains governed by *A*[*c*]. This distribution-independent result also admits a converse interpretation: for a fixed concentration-effect relationship, differences in baseline sensitivity heterogeneity alone are insufficient to reverse schedule ordering. An observed reversal across tumors or during treatment would therefore point to mechanisms beyond baseline distributional differences, such as changes in concentration-effect curvature, phenotype-specific response functions, phenotypic adaptation, or treatment-dependent sensitivity.

We assume that each phenotype has a fixed drug sensitivity *x* during treatment and that all phenotypes share the same concentration-effect function *f* (*c*), differing only through a non-negative sensitivity scale *x*. Under this assumptions, treatment can reweight the population by preferentially eliminating more sensitive phenotypes, but it does not alter the phenotype-specific response law itself. This assumption allows the effect of baseline heterogeneity and treatment-induced selection to be isolated from changes in pharmacodynamic shape or treatment-induced phenotypic adaptation. As a result, every phenotype remains coupled to the same schedule-dependent cumulative effect, making it possible to determine whether heterogeneity alone can reverse the schedule ordering established by concentration-effect curvature. The preservation result therefore applies specifically to selection among pre-existing phenotypes with a shared response shape.

Within this regime, these results caution against interpreting either post-treatment resistance or total drug exposure as direct measures of therapeutic efficacy. In our model, a schedule that generates a larger cumulative treatment effect can simultaneously produce greater tumor reduction and stronger enrichment for resistant phenotypes, because sensitive cells are preferentially removed. Thus, a more resistant surviving population does not necessarily indicate an inferior treatment; it may instead reflect more effective depletion of sensitive cells. A similar distinction applies to pharmacokinetic exposure. Quantities such as *C*_*max*_ and AUC summarize clinically relevant aspects of drug exposure, but equal exposure need not imply equal biological action when the concentration-effect relationship is nonlinear.^42^ Dynamic in vitro exposure studies have shown that the temporal distribution of drug concentration can alter treatment efficacy even when exposure is matched by conventional pharmacokinetic summaries.^39^ In our framework, this distinction is expressed mathematically: conventional exposure is represented by ∫ *c*(*t*)*dt*, whereas cumulative treatment effect is *A*[*c*] = ∫ *f* (*c*(*t*))*dt*. Therefore, the same total exposure can generate different biological effects depending on how that exposure is distributed over time. Together, these results show that neither survivor resistance nor total exposure is sufficient to characterize treatment efficacy.

Several extensions will be necessary to determine how robust this schedule-ordering principle remains in more dynamically evolving tumors. The current model assumes fixed phenotype-specific sensitivities and a shared concentration-effect function, thereby restricting treatment-induced change to selection among pre-existing phenotypes rather than phenotypic switching, adaptation, or evolution of the response function itself^11, 43–45^. Allowing sensitivity to become treatment-dependent, for example through *x*(*t*), an evolving distribution *p*(*x, t*), or phenotype-specific response function *f*_*x*_(*c*), could introduce genuine treatment-history dependence and potentially alter the ordering established here.

## Acknowledgments

The authors gratefully acknowledge funding by the National Cancer Institute via the Cancer Systems Biology Consortium (CSBC) U54CA274507; and Moffitt Cancer Center support from the Center of Excellence for Evolutionary Therapy.

## References

1. Scibilia, K. R. et al. Mathematical oncology: how modeling is transforming clinical decision-making. Cancer research 85, 4866–4879 (2025).

2. Altrock, P. M., Liu, L. L. & Michor, F. The mathematics of cancer: integrating quantitative models. Nature reviews cancer 15, 730–745 (2015).

3. Foo, J., Chmielecki, J., Pao, W. & Michor, F. Effects of pharmacokinetic processes and varied dosing schedules on the dynamics of acquired resistance to erlotinib in egfr-mutant lung cancer. Journal Thoracic Oncology 7, 1583–1593 (2012).

4. Köhn-Luque, A. et al. Phenotypic deconvolution in heterogeneous cancer cell populations using drug-screening data. Cell Reports Methods 3 (2023).

5. Pomeroy, A. E. & Palmer, A. C. A model of intratumor and interpatient heterogeneity explains clinical trials of curative combination therapy for lymphoma. Blood Cancer Discovery 6, 254–269 (2025).

6. Patterson, S. C., Pomeroy, A. E. & Palmer, A. C. Ultrasensitive response explains the benefit of combination chemotherapy despite drug antagonism. Molecular cancer therapeutics 23, 995–1009 (2024).

7. Skipper, H. E., Schabel, F. M. J. & Wilcox, W. S. Experimental evaluation of potential anticancer agents xiii, on the criteria and kinetics associated with” curability” of experimental leukemia. Cancer Chemotherapy Report 35, 3–111 (1964).

8. Skipper, H. E. The Effects of Chemotherapy on the Kinetics of Leukemic Cell Behavior. Cancer Research 25 (1965).

9. Perry, M. C. The Chemotherapy Source Book (Lippincott Williams & Wilkins, 2008).

10. Bozic, I. & Nowak, M. A. Resisting resistance. (2017).

11. Strobl, M., Gallaher, J., Robertson-Tessi, M., West, J. & Anderson, A. Treatment of evolving cancers will require dynamic decision support. Annals Oncology 34, 867–884 (2023).

12. Yu, H. et al. Phase 1 study of twice weekly pulse dose and daily low-dose erlotinib as initial treatment for patients with egfr-mutant lung cancers. Annals Oncology 28, 278–284 (2017).

13. Foo, J. & Michor, F. Evolution of resistance to targeted anti-cancer therapies during continuous and pulsed administration strategies. PLoS Computational Biology 5, e1000557 (2009).

14. Taleb, N. N. & West, J. Working with convex responses: Antifragility from finance to oncology. Entropy 25, 343 (2023).

15. Pierik, L., McDonald, P., Anderson, A. R. & West, J. Second-order effects of chemotherapy pharmaco-dynamics and pharmacokinetics on tumor regression and cachexia. Bulletin Mathematical Biology 86, 47 (2024).

16. Childers, L., Wiesch, P.A.z. & Conway, J. M. A general analytic approach to predicting the best antibiotic dosing regimen. Journal Biological Dynamics 20, 2641302, DOI: 10.1080/17513758.2026.2641302 (2026).

17. Harris, L. A. et al. An unbiased metric of antiproliferative drug effect in vitro. Nature Methods 13, 497–500 (2016).

18. Hafner, M., Niepel, M., Chung, M. & Sorger, P. K. Growth rate inhibition metrics correct for confounders in measuring sensitivity to cancer drugs. Nature methods 13, 521–527 (2016).

19. West, J. et al. Evolutionary antifragile therapy is a treatment strategy to suppress drug resistance by exploiting dose–response convexity. Cancer Research OF1–OF12 (2026).

20. Hare, L. et al. Drug holidays counteract alk inhibitor addiction in paediatric anaplastic large cell lymphoma. Leukemia Research 156 (2025).

21. West, J., Ma, Y. & Newton, P. K. Capitalizing on competition: An evolutionary model of competitive release in metastatic castration resistant prostate cancer treatment. Journal Theoretical Biology 455, 249–260 (2018).

22. Gallaher, J. A., Enriquez-Navas, P. M., Luddy, K. A., Gatenby, R. A. & Anderson, A. R. Spatial heterogeneity and evolutionary dynamics modulate time to recurrence in continuous and adaptive cancer therapies. Cancer Research 78, 2127–2139 (2018).

23. Hamis, S. et al. Growth rate-driven modelling suggests that phenotypic adaptation drives drug resistance in brafv600e-mutant melanoma. Communications Biology 9, 385 (2026).

24. Maley, C. C., Reid, B. J. & Forrest, S. Cancer prevention strategies that address the evolutionary dynamics of neoplastic cells: simulating benign cell boosters and selection for chemosensitivity. Cancer Epidemiology Prevention Biomarkers 13, 1375–1384 (2004).

25. Maley, C. C. et al. Classifying the evolutionary and ecological features of neoplasms. Nature Reviews Cancer 17, 605 (2017).

26. Qi, T., Dunlap, T. & Cao, Y. Embracing project optimus: Can we leverage evolutionary theory to optimize dosing in oncology? timothy et al. Pharmaceutical research 39, 3259–3265 (2022).

27. Gardner, S. N. A mechanistic, predictive model of dose-response curves for cell cycle phase-specific and-nonspecific drugs. Cancer research 60, 1417–1425 (2000).

28. Hill, B. T., Whelan, R. D., Rupniak, H. T., Dennis, L. Y. & Rosholt, M. A. A comparative assessment of the in vitro effects of drugs on cells by means of colony assays or flow microfluorimetry. Cancer Chemotherapy Pharmacology 7, 21–26 (1981).

29. Rupniak, H. T., Whelan, R. D. & Hill, B. T. Concentration and time-dependent inter-relationships for antitumour drug cytotoxicities against tumour cells in vitro. International Journal Cancer 32, 7–12 (1983).

30. Levasseur, L. M., Slocum, H. K., Rustum, Y. M. & Greco, W. R. Modeling of the time-dependency of in vitro drug cytotoxicity and resistance. Cancer Research 58, 5749–5761 (1998).

31. Keefe, D., Capizzi, R. L. & Rudnick, S. A. Methotrexate cytotoxicity for l5178y/asn-lymphoblasts: relationship of dose and duration of exposure to tumor cell viability. Cancer research 42, 1641–1645 (1982).

32. Slevin, M. et al. A randomized trial to evaluate the effect of schedule on the activity of etoposide in small-cell lung cancer. Journal Clinical Oncology 7, 1333–1340 (1989).

33. Clark, P. et al. A randomized trial of two etoposide schedules in small-cell lung cancer: the influence of pharmacokinetics on efficacy and toxicity. Journal clinical oncology 12, 1427–1435 (1994).

34. Scott, J. G. et al. A genome-based model for adjusting radiotherapy dose (gard): a retrospective, cohort-based study. The lancet oncology 18, 202–211 (2017).

35. Dale, R. Dose-rate effects in targeted radiotherapy. Physics Medicine & Biology 41, 1871–1884 (1996).

36. Nardi, V., Azam, M. & Daley, G. Q. Mechanisms and implications of imatinib resistance mutations in bcr-abl. Current opinion hematology 11, 35–43 (2004).

37. Taleb, N. N. (anti) fragility and convex responses in medicine. In International Conference on Complex Systems, 299–325 (Springer, 2018).

38. Axenie, C. et al. Antifragility in complex dynamical systems. npj Complexity 1, 12 (2024).

39. Komen, J. et al. Controlled pharmacokinetic anti-cancer drug concentration profiles lead to growth inhibition of colorectal cancer cells in a microfluidic device. Lab on a Chip 20, 3167–3178 (2020).

40. Bhang, H.-e. C. et al. Studying clonal dynamics in response to cancer therapy using high-complexity barcoding. Nature medicine 21, 440–448 (2015).

41. Patwardhan, G. A. et al. Treatment scheduling effects on the evolution of drug resistance in heteroge-neous cancer cell populations. NPJ Breast Cancer 7, 60 (2021).

42. Liston, D. R. & Davis, M. Clinically relevant concentrations of anticancer drugs: a guide for nonclinical studies. Clinical cancer research 23, 3489–3498 (2017).

43. Emond, R. et al. A novel combination therapy for er+ breast cancer suppresses drug resistance via an evolutionary double-bind. Molecular Systems Biology 1–27 (2026).

44. Vibishan, B. et al. Impacts of competition and phenotypic plasticity on the viability of adaptive therapy. Mathematical biosciences 109522 (2025).

45. Luddy, K. A. et al. Evolutionary double-bind treatment using radiotherapy and nk cell-based im-munotherapy in prostate cancer. International Journal Radiation Oncology* Biology* Physics (2025).

