## Supplementary for "Curvature-based schedule ordering is preserved in homogeneous and heterogeneous tumors"

### S1 Mathematical Proofs

We establish that the curvature-based ordering of equal-mean treatment schedules is preserved for any baseline distribution of fixed, nonnegative drug sensitivities.

From the population model in the main text, the final tumor burden under a concentration schedule $c(t)$ can be expressed as

$$N(T, c) = N_0\Phi(T)E_{p_0}[e^{-XA[c]}] = N_0\Phi(T)\mathcal{L}_{p_0}(A[c])$$

where $\mathcal{L}_{p_0}$ denotes the Laplace transform of the baseline sensitivity distribution, evaluated at $A[c]$.

For a continuous distribution, the Laplace transform is defined by

$$\mathcal{L}_{p_0}(A) = E_{p_0}[e^{-XA}] = \int_0^\infty p_0(x)e^{-xA}dx, \qquad A \geq 0.$$

We first show that a larger cumulative treatment effect cannot produce a greater final tumor burden, regardless of the baseline sensitivity distribution.

*Proof.* Since $x \geq 0$, $A[c_1] \geq A[c_2]$ implies $xA[c_1] \geq xA[c_2]$ for $\forall x \geq 0$. Because $e^{-xA}$ decreases monotonically with $A$, $\forall x \geq 0$. Multiplying by $p_0(x) \geq 0$, integrating over $x$, and multiplying by $N_0\Phi(T) > 0$ gives $N(T, c_1) \leq N(T, c_2)$. □

Next, we consider an even schedule $c_e(t) = \bar{c}$ and uneven schedule $c_u(t)$ with the same temporal meanl. Jensen's inequality determines the ordering of their cumulative treatment effects, which in turn determines the ordering of final tumor burden.

*Proof.* For concave $f$, Jensen's inequality gives $\frac{1}{T}\int_0^T f(c_u(t))dt \leq f(\bar{c})$, i.e. $A[c_u] \leq Tf(\bar{c}) = A[c_e]$. For convex $f$, Jensen's inequality reverses, giving $A[c_u] \geq A[c_e]$, and therefore, $N(T, c_u) \leq N(T, c_e)$. □

For the power-law response $f(c) = c^h$, even exposure is favored when $0 < h < 1$, whereas uneven exposure is favored when $h > 1$. At $h = 1$, both schedules yield identical final tumor burdens. The ordering is strict when the cumulative treatment effects differ and the baseline population assigns positive probability to $X > 0$.

We show that treatment-induced selection shifts the surviving population toward lower drug sensitivity, while the final tumor burden remains determined by the total cumulative treatment effect.

From the survivor sensitivity distribution defined in the main text, the mean sensitivity among surviving cells is

$$\bar{x}(A) = \frac{E[Xe^{-XA}]}{E[e^{-XA}]}.$$

*Proof.* Differentiating with respect to the cumulative treatment effect A using the quotient rule gives

$$\frac{d\bar{x}}{dA} = -\frac{E_{p_0}[X^2 e^{-XA}]}{E_{p_0}[e^{-XA}]} + \bar{x}(A)^2 = -Var_A(X). \leq 0$$

Thus, the mean sensitivity of surviving cells is non increasing in cumulative treatment effect. □

### S2 Numerical Implementation

Numerical simulations were performed in Python using NumPy, with figures generated using Matplotlib. Because the population model admits an exact exponential solution for prescribed concentration trajectories, tumor burden and survivor sensitivity distributions were evaluated directly without numerically solving the governing differential equation. For the power-law concentration-effect function $f(c) = c^h$, cumulative treatment effects were calculated analytically for even and first-order pharmacokinetic (PK) schedules, by exact piecewise integration for step schedules, and by cumulative trapezoidal integration for sinusoidal schedules. Integrals over continuous sensitivity distributions were evaluated using the trapezoidal rule.

Equal-mean concentration schedules were constructed over dosing cycles of duration $\tau$, with mean concentration $\bar{c} = 1$. The step schedule used an on-fraction of 0.5, alternating between concentrations of 2 and 0. The sinusoidal schedule had amplitude 1, while the periodically repeated first-order PK schedule used an elimination rate of 1.5 and an initial concentration of approximately 4.55055 to satisfy the equal-mean condition. Continuous baseline sensitivity distributions were evaluated on finite numerical grids and normalized to unit probability mass. Survivor distributions were obtained by exponential reweighting of the baseline density followed by normalization, and mean survivor sensitivity was calculated by numerical quadrature.

For the distributional robustness analysis (Fig. 5), lognormal, gamma, and Weibull distributions were parameterized to have a common mean of 1 and variance of approximately 0.1929. Gamma parameters were determined from the target moments, and the Weibull shape parameter was obtained numerically by matching the coefficient of variation.

### S3 Code Availability

The Python code used to generate the numerical illustrations in Figures 1-5 is publicly available at [https://github.com/integralong/curvature-schedule-ordering]. All numerical results were obtained from model-based simulations using synthetic parameter settings; no empirical biological datasets were used.
